# Propidium monoazide (PMA) with quantitative PCR to detect and quantify viable *Paenibacillus larvae* and *Melissococcus plutonius* in honeybee hive samples

**DOI:** 10.64898/2026.02.20.707117

**Authors:** Anuja Shrestha, Brandon Kingsley Hopkins, James T. Van Leuven

**Affiliations:** Department of Animal Veterinary and Food Science, University of Idaho, Moscow, ID, USA; Department of Entomology, Washington State University, Pullman, WA, USA

**Keywords:** *Paenibacillus larvae*, *Melissococcus plutonius*, propidium monoazide, quantitative PCR, viable, ozone

## Abstract

*Paenibacillus larvae* and *Melissococcus plutonius* are the bacterial agents responsible for American and European foulbrood diseases, respectively, and pose significant economic threats to apiculture worldwide. Early detection of these pathogens is critical for effective disease prevention and management in honeybee colonies. *P. larvae* forms highly persistent spores that can remain infectious on hive equipment for decades, whereas the non-spore-forming *M. plutonius* can survive for over a year in contaminated materials. Although various decontamination and treatment strategies are available or under development, reliable methods for quantifying these pathogens are essential for accurately assessing their effectiveness. Culture-based techniques and quantitative PCR (qPCR) are commonly used to detect and quantify foulbrood pathogens in hive-associated samples; however, culturing is labor-intensive and sometimes inconsistent, while qPCR does not distinguish between viable and dead cells. Propidium monoazide (PMA) selectively binds DNA from dead cells and prevents its amplification during qPCR, enabling viable-cell quantification. Despite this advantage, the performance of PMA-qPCR for detecting *P. larvae* and *M. plutonius* in hive samples has not previously been assessed. In this study, we evaluated PMA-qPCR using both pure cultures and environmentally relevant honeycomb samples before and after ozone treatment. PMA treatment greatly reduced the erroneous detection of dead cells by several orders of magnitude in both culture and hive samples, supporting its utility for AFB and EFB risk assessment and treatment evaluation. However, incomplete suppression of dead-cell DNA at high cell densities indicates that further optimization of PMA treatment conditions may be necessary.

## INTRODUCTION

Honeybees (*Apis mellifera* L.) play a major role in crop pollination, directly facilitating the enhancement of crop yield and quality, and contributing approximately one-third of the total human food supply (Khalifa et al., 2021). In the United States alone, honeybees pollinate $15 billion worth of crops annually (Morse & Calderone, 2000; Kulhanek et al., 2017). Commercial beekeepers transport more than two million colonies around the United States annually, to pollinate crops like almonds, apples, blueberries, and melons (Bond et al., 2021; Goodrich, 2019), a practice called ‘pollination migration’. The role of honeybees as crop pollinators is at risk due to multiple biotic and abiotic stressors, which cause widespread colony losses and lead to significant economic losses (French et al., 2024). Among the multiple stressors, infectious diseases such as American foulbrood (AFB) and European foulbrood (EFB) are severe bacterial diseases that primarily affect honeybee larvae, causing substantial economic losses in apiculture worldwide (Forsgren et al., 2010; Genersch, 2010).

AFB is caused by a gram-positive, spore-forming bacterium, *Paenibacillus larvae*, which is highly infectious and can kill the infected bee colonies (Genersch et al., 2005; Kleijn et al., 2015). The larvae get infected by the ingestion of spore-contaminated food, which germinates and proliferates in the midgut, invades the larval tissue, and kills the host larvae (Ghorbani-Nezami et al., 2015; Yue et al., 2008). The decayed larvae host millions of spores, which can be transmitted to larvae by nurse bees or by spores remaining in the brood cells. In addition, exchanging diseased brood combs, feeding on or robbing infected honey or bee bread, trading queens and colonies, and producing wax foundations contaminated with spores are other potential means of spore transmission within and between colonies (Lindström, 2008). Furthermore, these spores are extremely heat-stable, resistant to disinfection, and can remain infectious for several decades in honey or on hive equipment, making the effective control of AFB difficult (Genersch, 2010; Morrissey et al., 2015). Molecular techniques have confirmed five genotypes of *P. larvae*: ERIC I and ERIC II are the most commonly isolated genotypes from infected hives (Matović et al., 2023); ERIC III and IV are more lethal; however, they have not been isolated in the field in decades (Ebeling et al., 2016), and ERIC V genotypes were recently discovered (Beims et al., 2020). The infection and transmission methods of these genotypes are similar, whereas the notable difference is virulence (Rauch et al., 2009). Hives infected with AFB require intervention and are a significant cause of economic loss for the beekeeping industry (Genersch, 2010; Kušar et al., 2021); addressing this disease is also a legal requirement in most European countries (Genersch, 2010; Locke et al., 2019). Prevention and treatment measures include the use of antibiotics (Masood et al., 2022), breeding for honeybee immune response against AFB (Rauch et al., 2009; Spivak & Reuter, 2001), biocontrol using antagonistic bacteria (D Evans & Armstrong, 2005), treatment with natural antibacterial substances like essential oils and plant extracts (Ebert et al., 2007), decontaminating hive materials, etc. The most effective way to prevent AFB spread is to detect the disease early and destroy contaminated hive materials and often the colony itself. General symptoms of AFB are irregular brood capping, with capped and uncapped cells scattered irregularly across the brood frames. Infected cells become dark, sunken, and emit a foul odor. The brown, glue-like remains of the dead larvae in these cells form a highly characteristic ropey thread when pulled out with a wooden stick or a match, which is a reliable field diagnosis for AFB. The remains of these larvae form a hard scale at the bottom of the cells, containing millions of spores (Forsgren & Laugen, 2014).

EFB is caused by the gram-positive, non-spore-forming bacterium, *Melissococcus plutonius*, which causes an intestinal infection of honeybee larvae and is transmitted by symptomless adult honeybees to young larvae via contaminated food (Takamatsu et al., 2015). The infectious cycle begins when larvae ingest brood food contaminated with *M. plutonius*. The bacterium then reaches the midgut, localizes there, and multiplies in the midgut lumen and the peritrophic matrix that lines the midgut epithelium, gradually filling it, competing with the host for nutrients, and leading to larval death through starvation (Bailey, 1983; Forsgren, 2010). The massive loss of brood from severe infection weakens the colony and can even lead to its collapse (Grossar et al., 2020). *M. plutonius* consists of two groups of strains: typical and atypical strains, based on multilocus sequence typing (MLST), which are phenotypically and genetically distinguishable (Arai et al., 2012). These strains are further divided into three genetically distinct clonal complex groups based on the sequence types (STs) by MSLT: atypical strains are grouped into CC12, and typical strains into CC3 and CC13 (Haynes et al., 2013; Takamatsu et al., 2014). EFB disease has been reported worldwide and is also classified as a notifiable disease because of the severity of EFB outbreaks and the absence of efficient treatments (Grossar et al., 2020). Unlike AFB, hives can also spontaneously recover from EFB on their own due to the high variability in the pathogen’s virulence (Milbrath et al., 2021; Mosca et al., 2024). EFB tends to be more prevalent under conditions of stress, such as locations with nectar or pollen dearths. Different control and management methods are recommended, such as the destruction of the colony, shook swarm techniques, antibiotic treatment (Waite et al., 2003), and integrated pest management, including good hive hygiene, proper nutrition supplementation, replacing the queen, and replacing old equipment to prevent pathogen buildup (Mosca et al., 2024). Contaminated equipment can harbor the bacteria for a year or more, even without a spore stage (Wine, 2021), but it is considered less severe and treatable compared to spore-forming AFB.

Decontaminating hive material can be done by chemical disinfection, scorching with a blowtorch, immersion into molten paraffin, methyl oxide & gamma radiation, ozone treatment, etc. (Torlak & Işik, 2018). These treatments have their advantages as well as drawbacks. The use of ozone as a sanitizer is known to be an environmentally friendly treatment, as it has been registered for direct application on the surface by the US Environmental Protection Agency (Achen & Yousef, 2001; Guzel-Seydim et al., 2004). It has also been reported to be more practical as compared to other sanitizers, as it doesn’t require storage, special handling, or mixing considerations (Torlak & Işik, 2018). It can disinfect and eliminate odors, taste, and color, and has also been used as an agricultural fumigant to kill insects and pests in stored grains (James, 2011; Mendez et al., 2003). However, it is important to check if any of these decontamination/control methods can actually eliminate pathogens from hive materials.

Detection methods for AFB and EFB include apiary inspection for visual signs and symptoms, and are often confirmed with further testing using commercial field test kits and/or laboratory diagnosis, such as microscopic examination, lateral-flow devices, culture-based methods, and molecular techniques such as PCR (Heo et al., 2025). Visual inspection of the brood frame provides subjective interpretations of clinical signs of infections, which often require additional confirmation. Microscopic examination of larval smears is insufficient for reliable detection of bacterial loads. Culture-based methods are widely used for detecting and quantifying *P. larvae* and *M. plutonius* in samples associated with beehives (Papić et al., 2024). However, quantifying cells through plate counting is time-consuming, sometimes unreliable due to poor and inconsistent germination, and requires species confirmation (Kušar et al., 2021). Molecular quantification techniques, such as PCR and qPCR, enable rapid detection and quantification of target DNA with high sensitivity and specificity, including viable but non-culturable cells. However, these methods amplify all DNA from both viable and dead bacteria, making it impossible to differentiate between them (Okada et al., 2022; Rodgers et al., 2017). To address this issue, several studies have utilized propidium monoazide (PMA) in combination with PCR or qPCR to detect and quantify viable cells in various non-apiary samples (Fu et al., 2020; Heise et al., 2016; Nocker et al., 2007; Rogers et al., 2010). PMA is a DNA-intercalating dye that selectively enters compromised cells through the damaged cell membrane and binds to their DNA, thereby inhibiting PCR amplification and allowing for the distinction between live and dead cells (Nocker et al., 2007). PMAxx™, a modified version of PMA developed by Biotium, Inc. (Fremont, CA, United States), has been reported to enhance the reactivity of PMA.

To the best of our knowledge, no published studies have used PMA or PMAxx™ with qPCR to detect or quantify viable *P. larvae* and *M. plutonius* in beehive samples. This presents a promising approach to detect the pathogens as well as to evaluate the effectiveness of different control measures tested against AFB and EFB diseases. Therefore, we hypothesized that PMAxx™, in conjunction with SYBR Green-based quantitative PCR, could accurately and consistently detect and quantify viable *P. larvae* and *M. plutonius* in hive samples, so that better and more reliable risk assessment and treatment options can be tested and applied. We also hypothesized that ozone treatments are lethal and could disinfect the *P. larvae* pathogen from the AFB-infected bee hives, although evaluating the effectiveness of ozone was not the purpose of this study.

## MATERIALS AND METHODS

### Bacterial strains and culture conditions for *Paenibacillus larvae* strains

Two *Paenibacillus* strains, *P. larvae* subsp *larvae* strain Y-3650 (NRRL B-3650) (ERIC group I), and *P. larvae* subsp *pulvifaciens* strain 368 (ATCC-25368) (ERIC group IV), were used. Y-3650 was obtained from the University of Nevada, Las Vegas (Yost et al., 2016), and 368 from the American Type Culture Collection (ATCC) Genbank. These strains were retrieved by streaking glycerol stocks stored at −80°C freezer onto 1mg/L of thiamine hydrochloride Brain Heart Infusion (BHI) agar (Catalog#DF0418-17-7, BD DifcoTM, USA) plates and incubated at 37°C with 5% CO2 for 72 hours. A single colony from the plates was suspended in 3 mL of BHI broth (Catalog#CM1135B, Thermo Scientific™, USA) and incubated for 24 hours at 37°C with shaking at 120 rpm using a MaxQ™ 6000 incubator shaker (Catalog SHKE6000, Thermo Scientific™, USA). One mL of the cultures was suspended in 9 mL of BHI broth in sterile flasks to make subcultures and then incubated for 24 hours at the abovementioned conditions. The optical density of the liquid cultures in flasks at 600 nm was adjusted to 1.0 using a SpectraMax iD3 Microplate Reader (Molecular Devices, CA, USA), which corresponded to a final concentration of approximately 10^8^ CFU/mL for both strains.

### Bacterial strains and culture conditions for *Melissococcus plutonius* strains

Two *M. putonius* strains, ATCC-35311 (typical strain, ST-1, CC13) (Djukic et al., 2018; Grossar et al., 2023; Haynes et al., 2013) and 19-18-9 (atypical strain, ST-12, CC-12) were used. Strain 19-18-9 was obtained from Michigan State University, Michigan, and ATCC-35311 from ATCC GenBank. Both bacterial strains were streaked on freshly prepared M110 agar media (Budge et al., 2024), and plates were incubated in an anaerobic chamber with 5% CO_2_, 10% H_2_, and 85% N_2_ at 37 °C for 3-6 days. For strain 19-18-9, colonies were used to prepare liquid cultures in KSBHI (brain heart infusion (BHI) supplemented with 0.15 M KH_2_PO_4_ and 1% soluble starch) and incubated anaerobically at 37 °C for 3-4 days at 150 rpm. For the ATCC-35311 strain, the cultures were prepared in ATCC 1430 broth (ATCC; Garrido-Bailon et al., 2013) and incubated at 30 °C for 4 days at 150 rpm. The bacterial pellets were collected by centrifuging the suspensions at 10,000 rpm for 10 minutes, and the supernatants were completely removed. The collected cell pellets were resuspended in their respective broth, and the optical density at 600 nm was adjusted to 1, as mentioned earlier, corresponding to the final cell concentrations of 10^6^ CFU/mL for both strains.

### Preparation of viable and dead cells of *Paenibacillus larvae* and *Melissococcus plutonius* strains

The bacterial suspensions of various concentrations were prepared by serially diluting the 10^8^ CFU/mL cell cultures of the *P. larvae* strains Y-3650 and 368 in PBS. Three different cell dilutions of each strain, with cell concentrations adjusted to approximately 10^8^, 10^6^, and 10^4^ CFU/mL, were used as the source of viable cells. Dead cells of each strain were generated by heating all cell dilutions at 95 °C for 20 minutes using a standard laboratory heat block. Cell viability was examined by plating 100 μL of each dilution on BHI plates and incubating the plates for 3 days. No cell growth in all plates means cells were dead in all dilution plates. However, there was cell growth on the 10^^8^ and 10^^6^ cell concentration plates of strain 368. Therefore, all three cell dilution cultures of 368 were exposed to 120°C for 20 minutes and plated again, making sure there was no cell growth on the incubated plates.

### M. plutonius strains

The 10^6^ CFU/mL cell cultures of both strains were serially diluted, and cell concentrations of 10^6^, 10^4^, and 10^2^ CFU/mL were used for viable cells. For dead cells, these dilutions were heat-treated at 95 °C for 30 minutes using a water bath, followed by plating onto freshly prepared M110 agar plates and incubating for 3-6 days in an anaerobic chamber to assess cell viability.

### AFB-infected comb sample collection and ozone treatment

Honeycomb samples infected with AFB were collected from Pullman, Washington, USA. Both microbiome and culture-based detection methods were used to confirm the presence of *P. larvae* in the samples. For the microbiome study, DNA was extracted from the samples using the ZymoBIOMICS DNA Microprep Kit, following the manufacturer’s protocol, and the DNA samples were sent to SeqCenter, Pittsburgh, PA, for 16S Illumina sequencing. For the culture-based study, the samples were processed and cultured in our lab (Shrestha et al., 2025). In short, the samples were ground with the phosphate buffer, serially diluted, and plated onto BHI agar, incubated for 3 days, purified unique colonies, made cultures in BHI broth, and extracted DNA. The extracted DNA samples were sent to Eurofin Genomics, DNA Sequencing Lab, Louisville, USA, for 16S rRNA sequencing. Both these detection methods confirmed the presence of *P. larvae* in the samples. The AFB-infected comb samples were divided into 4 groups: two used as controls (untreated with ozone) and two were ozone-treated. For the ozone treatments, the samples were kept in an ozone chamber for an hour at a concentration of 1000 parts per million (ppm). The ozone chamber was set at 30 °C temperature at WSU, Puyallup Campus, Washington, USA. After the treatments, swabs from the comb samples were collected from both untreated (controls) and ozone-treated samples and stored in 1.5 mL tubes with 500 µL of 40% glycerol.

### PMAxx™ treatment

For PMA treatment, we used cell concentrations of 10^8^, 10^6^, and 10^4^ CFU/mL for *P. larvae* strains and 10^6^, 10^4^, and 10^2^ CFU/mL for *M. plutonius* strains. Two light-transparent 1.5 mL microcentrifuge tubes, each with 400 μL of either live cell or dead cell cultures or swab samples from each treatment (controls and ozone-treated), were used: one tube for PMAxx™ treatment and the other for no PMAxx™ treatment. Under dark conditions, PMAxx™ solution (20 mM in H_2_O; Biotium Inc.) was added to 400 μL of the samples to achieve a final concentration of 25 μM. The tubes were incubated for 10 minutes in a rocker set up at room temperature in the dark for optimum mixing. The photoactivation of tubes was done by exposing them to light (12,000 Lumens LED lamp, BRAUN, China) for 20 minutes in a rocker, which helps the PMAxx™ dye to covalently bind to the dead cell DNA (Figure 2). To minimize the effect of light radiation, the microcentrifuge tubes were placed horizontally on ice, and the distance between the tubes and the lamp was maintained at around 20 cm. The tubes with the live cell or dead cell cultures were centrifuged at 5,000 x g for 10 minutes to collect cell pellets for further DNA extraction. In contrast, the swab samples were proceeded to DNA extraction without centrifugation.

### DNA extraction and quantitative PCR analyses

DNA was extracted using a Quick-DNA HMW MagBead Kit (Zymo Research, Catalog # D6060), following the manufacturer’s protocol. The final elution volume was 50 μL. The concentration of the extracted DNA was measured by NanoDrop™ One (Catalog# ND-ONE-W, ThermoFisher Scientific, United States). The extracted DNA without normalizing the DNA concentration was used for quantitative PCR (qPCR) analysis using QuantStudio™ Real-Time thermal cycler (Applied Biosystems, ViiA 7 by Life Technologies, USA). The PCR was carried out in 10 μL reactions with the PowerUp SYBR Green Master Mix (Applied Biosystems, ThermoFisher Scientific, USA). For *P. larvae* strains, 2 μL of DNA, 5 μL Master mix, 0.5 μL of PLAup forward primer, 0.5 μL of PLAdw reverse primer (Rossi et al., 2018), and nuclease-free water were added to reach the reaction volume. The PCR program consisted of an initial denaturation at 95 °C for 4 minutes and 40 cycles of denaturation at 95 °C for 15 seconds and annealing at 56 °C for 30 seconds, followed by melting curve analysis. For *M. plutonius*, different primers were used (Table 1): for the typical strain, Mp-Trt-F and Mp-Trt-R primers, and for the atypical strain, Mp-Art-F and Mp-Art-R were used (Nakamura et al., 2016). The amplification cycle consisted of initial denaturation at 95 °C for 15 minutes and 45 cycles of denaturation at 95 °C for 15 seconds, annealing at 53 °C for 30 seconds, and extension at 72 °C for 30 seconds. Each sample was run in duplicates, with nuclease-free water used as a negative control.

**Table 1.**
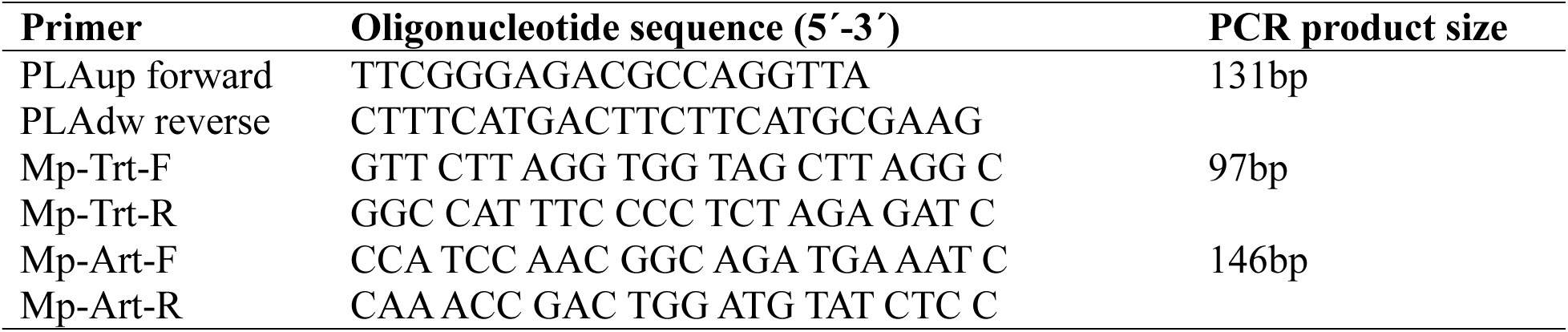
Primers used in this study. PLA primers specifically amplify *P. larvae*. Mp-Trt primers specifically amplify typical *M. plutonius*. Mp-Art specifically amplifies atypical *M. plutonius*.

All statistical analyses were performed in R. Bacterial loads between different groups were compared using a paired t-test. The value P ≤ 0.05 was considered the threshold for significance. The regression slope for live-cell, untreated samples was −10.0 (p = 1.25E-4, linear model, Ct ~ cell concentration + treatment group). This regression suggested that primer efficiency was good. For other treatments (e.g., PMA-treated, dead cells) with higher Ct values, we no longer observed this perfect linear relationship. Thus, we calculated the percent error (Expected Ct – Observed Ct / Expected Ct) * 100, where the expected Ct was based on the dilution amount, and found high error rates for samples with Ct values above ~35. Thus, when calculating the copy number difference of PMA-treated to untreated samples, we only used the highest cell dilutions.

## RESULTS

### Testing conditions required to kill the honeybee pathogens, *Paenibacillus larvae* and Melissococcus plutonius

To generate dead cells for testing PMA, we tested various heat-killing treatments to kill *P. larvae* and *M. plutonius*. We found that *P. larvae* strain 368 is more durable than the other strains, surviving boiling conditions for 20 minutes (Figure 1, Table 2). Because of this, we increased the temperature of heat treatment for strain 368 to 120°C. Strain Y-3650 was killed by the 95°C treatment. *M. plutonius* was tested at 95°C for 30 min, which was sufficient to not recover any live cells. The heat-killed dead cells were confirmed by not observing any colonies on agar plates (see details in Materials and Methods Sections).

**Figure 1.**
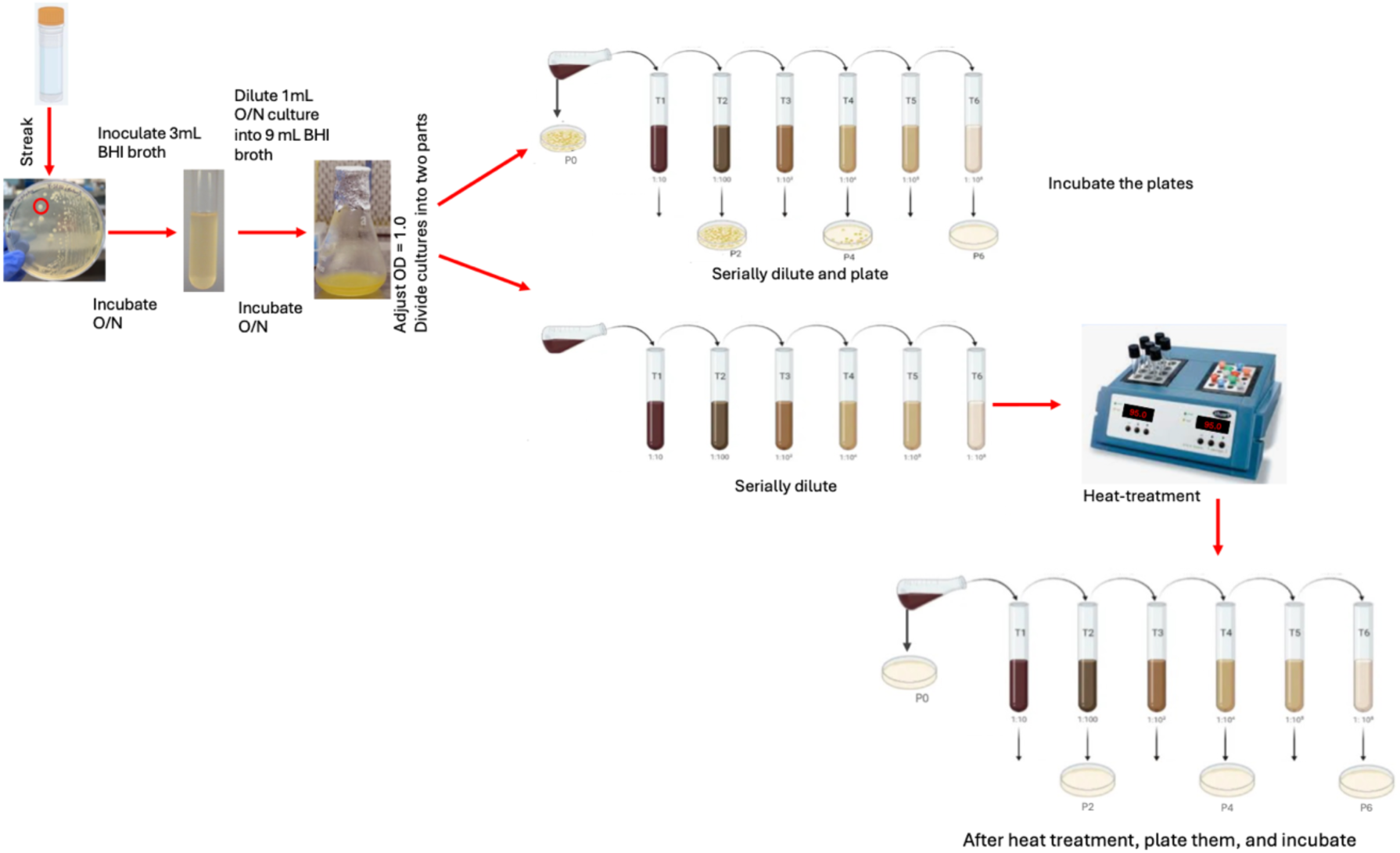
Diagram showing the steps of producing live and dead cells from a bacterial culture. Glycerol stocks of stored bacteria were plated on Brain Heart Infusion (BHI) agar plates (see methods) and incubated overnight (O/N). Single colonies were chosen for liquid growth. The optical density (OD) of liquid cultures at 600 nanometers was used to approximate cellular concentrations. Agar plates were used to accurately measure cell density (P0) before dilution series were made.

**Table 2.** Testing conditions required to kill bacterial pathogens. The cell concentrations before and after the listed treatments are shown.

| Strains | Live cells | Heat-kill temperature at |  |  |
| --- | --- | --- | --- | --- |
|  | CFU/mL | 95°C for 20 min | 120°C for 20 min | 95°C for 30 min |
| Y-3650 | 5.0 X 10<sup>8</sup> | 0 | n.a | n.a |
| 368 | 1.3 X 10<sup>8</sup> | 1.7 X 10<sup>5</sup> | 0 | n.a |
| ATCC-35311 | 5.7 X 10<sup>6</sup> | n.a | n.a | 0 |
| 19-18-9 | 8.2 X 10<sup>6</sup> | n.a | n.a | 0 |

n.a= not applicable (not tested)

### Testing PMA-qPCR with heat-killed cultures of *Paenibacillus larvae* and *Melissococcus plutonius*

Using the same samples from the above culture work, we made 100-fold and 10,000-fold dilutions of *P. larvae* and *M. plutonius* cell cultures, treated with PMAxx, extracted DNA, and then quantified DNA using qPCR (see methods for details, Figure 2). The addition of PMAxx reduced the detection of dead pathogens by qPCR across all strains and dilutions by an average of 31,723-fold (p=1.6E-7, t-test). This large difference was not observed with live cells (Figure 3), where we saw only a 166-fold increase in genome copy number detected without versus with PMA (p=0.025, t-test). Given that our null expectation was that live cells would entirely prevent PMA from binding to genomes, we were surprised to see this difference in live cells. Upon further investigation, we found that the amplification for *M. plutonius* strain 19-18-9 was potentially problematic, with high Ct values. When this strain is removed, the addition of PMA to live cells only changes genome quantification by 5-fold instead of 166-fold. For all strains except for 19-18-9, we were able to use the 100-fold dilutions to calculate the deviation from expected Ct values using 2^△Ct^ (see methods for details). This calculation confirmed that samples with Ct values above around 35-37 had large deviations from expected based on the dilution (Supplemental Figures 1 and 2). We used the atypical primers (Table 1) for strain 19-18-9 and found only dim bands when visualized on an agarose gel (Supplemental Figure 3). These two pieces of evidence suggest that while the atypical primers do amplify 19-18-9, they do so with poor efficiency, despite thermocycling conditions identical to published reports. Leaving out strain 19-18-9, we observe a significant difference between the effect of PMA on heat-killed versus live cells for every strain and dilution (Figure 3), although the relative reduction is impacted by cell concentration. As we used the suggested PMAxx concentration and incubation times, further optimization could result in more uniform inhibition. The cell densities that we used in the test studies ranged from 1×10^^2^ to 1×10^^8^ cells/mL, which likely encompasses the realistic cell densities of pathogens in bee larvae, but probably higher than those on hive surfaces such as tools or boxes. Another observation to note is that at high cell densities, we had amplification of heat-killed cell cultures that were treated with PMAxx. No colonies were observed when these same samples were plated on agar. This further suggests that optimization of PMA treatment could improve our results. However, just using the manufacturer’s suggested PMA conditions and previously reported PCR conditions, adopting this method could reduce the overestimate of bacterial pathogens in a sample by several logs.

**Figure 2.**
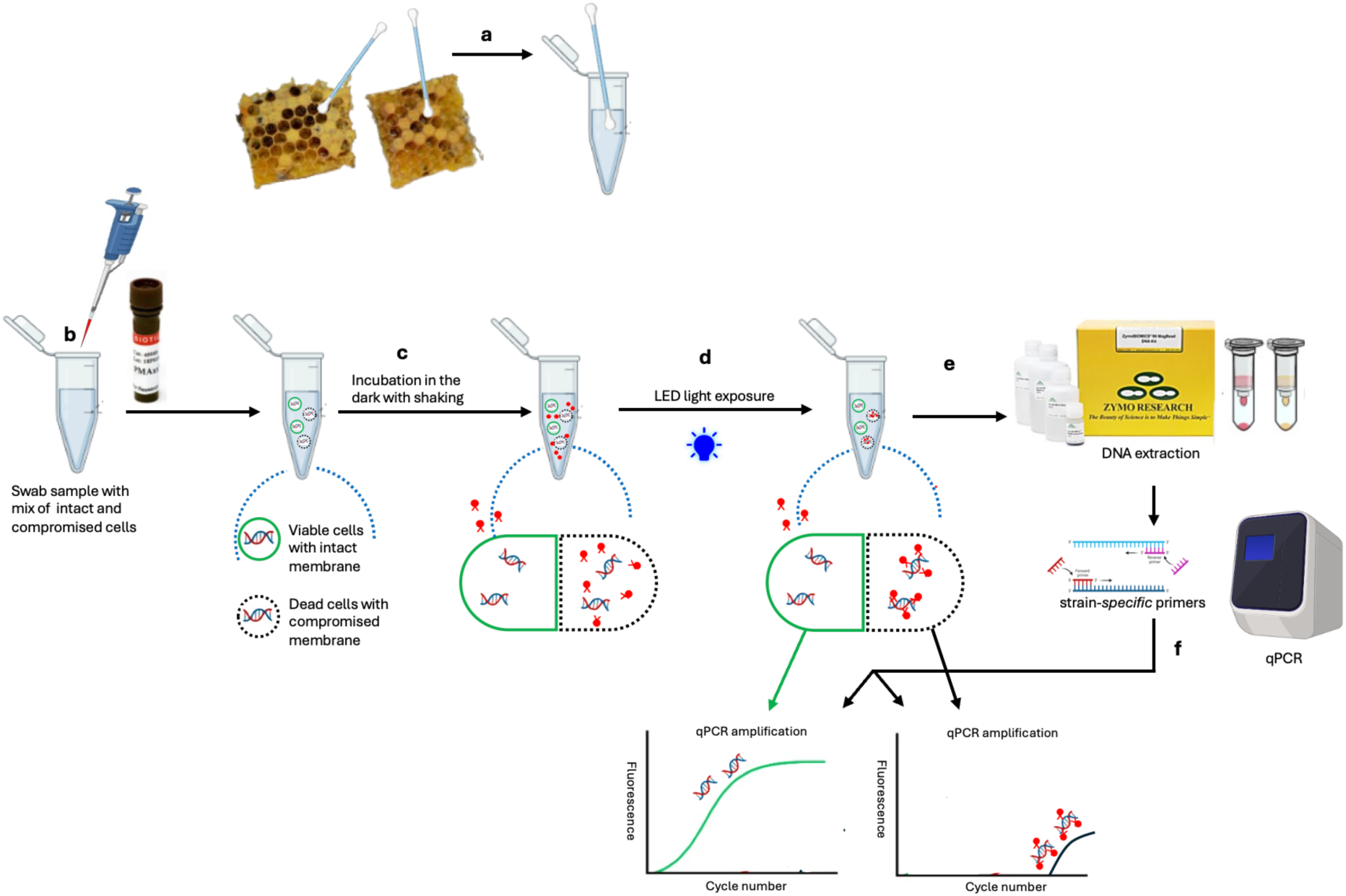
Methodology and mechanism of action of PMA-qPCR reaction: (a) swab collection; (b) addition of PMAxx dye; (c) incubation in the dark with shaking for optimal mixing between DNA and PMAxx dye; (d) photoactivation step to cross-link PMAxx to DNA, the PMAxx covalently bind to the dead cell DNA; (e) DNA extraction from the swab samples; (f) perform qPCR for the amplification of the target sequence.

**Figure 3.**
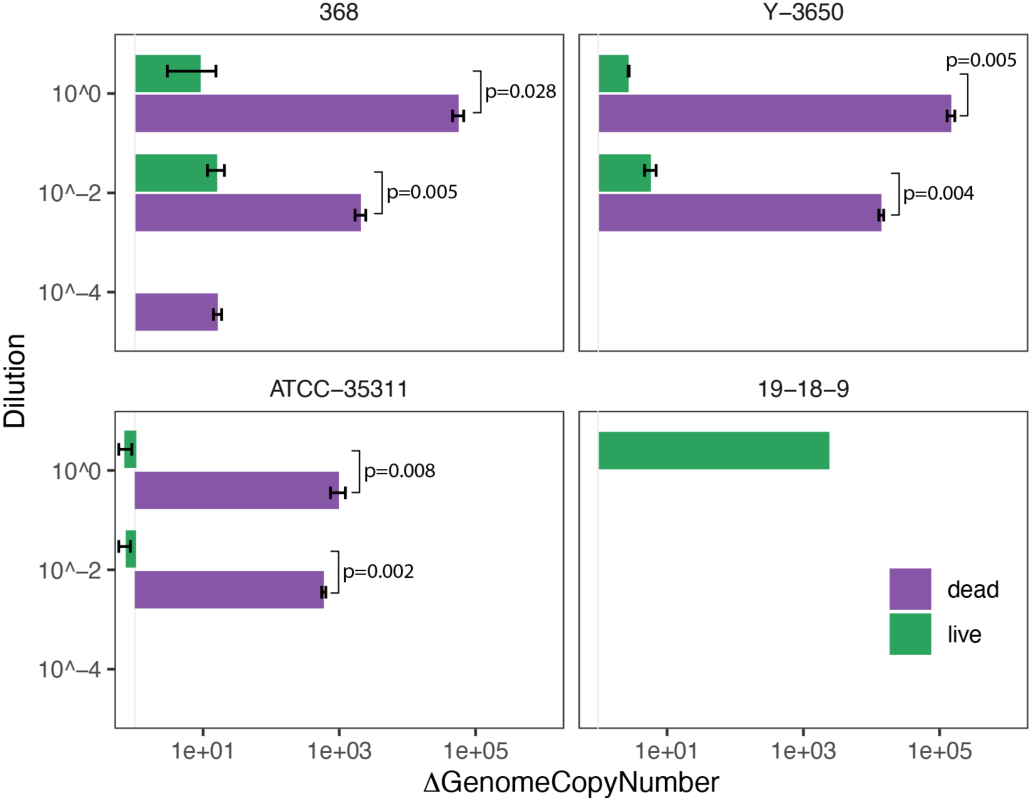
PMA treatment has a large impact on the quantification of dead *Paenibacillus larvae* and *Melissococcus plutonius* cells. The difference in the genome copy number (ΔGenome Copy Number) between PMAxx-treated and untreated samples is shown for heat-killed (dead) and live pathogens from cell culture at three dilutions (different cell densities). Error bars represent one standard deviation. Significant p-values show the differential effect of PMA between live and dead (heat-killed) cells.

### Impact of PMA treatment on *Paenibacillus larvae* quantification in ozone-treated vs non-treated honeycomb samples from AFB-infected hives

Given how well PMA prevents amplification of DNA from dead cells in culture, we tested the method on a more realistic environmental sample. We used *P. larvae*-infected honeycomb samples from a separate ozone treatment experiment. The comb was exposed to ozone for an hour while measuring the viability of *P. larvae*. After washing the comb with PBS, we split the samples in half for treatment with PMA. The cell density in these samples was measured at 4.0 × 10^5 CFU/mL by plate agar. *P. larvae* was then quantified using PMA-qPCR. In comb samples before ozone treatment, we observed 131 times fewer genome copies in the PMA-treated comb compared to the washed comb without PMA treatment (p = 0.002, paired t-test).

After one hour of ozone exposure, this difference was reduced to only 9.9-fold (p=0.017, paired t-test) (Figure 4). A possible reason for this decrease to the degradation of extracellular DNA during exposure to ozone. It is worth noting that both PMA-treated and untreated samples had lower *P. larvae* genome copies detected after ozone treatment, but the difference between PMA and no-PMA was significantly lower after ozone treatment (Figure 4). This result shows that PMA may be used on hive samples to differentiate between live and dead pathogens.

**Figure 4.**
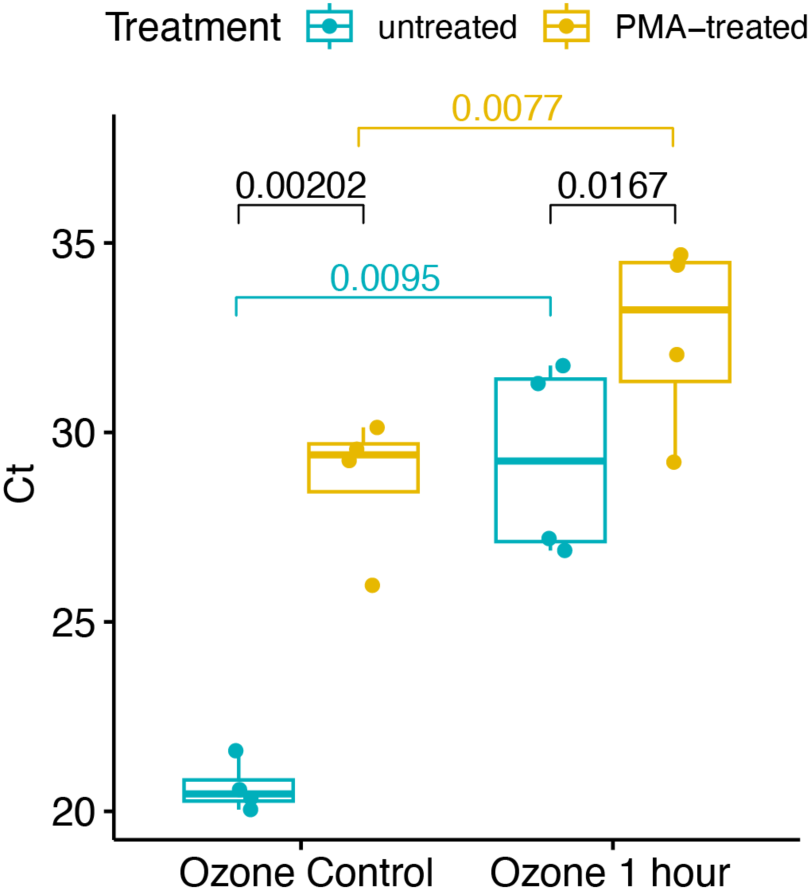
PMA-qPCR reduces the detection of dead *P. larvae* cells from comb samples. Fluorescent signal (Ct) of PMA-treated vs. untreated comb samples were compared prior to and after 1 hour of ozone treatment.

## DISCUSSION

Due to the high transmissibility of AFB and its regulatory consequences, timely and precise diagnosis is critical to prevent colony losses and limit further spread (Ebeling et al., 2016). In contrast, EFB is less likely to be transmitted and is often treated with antibiotics (Waite et al., 2003); however, the routine use of antimicrobials may promote the development of antimicrobial-resistant strains, thereby reducing treatment effectiveness (Reybroeck et al., 2012). Therefore, early and accurate detection is essential. To date, inspections for visual symptoms, confirmed using commercial test kits and laboratory diagnostic methods such as microscopic examination, PCR testing, or cultivation techniques, are commonly employed for disease detection. However, these methods do not detect and quantify viable pathogens. The detection of foulbrood pathogens from apiary samples using a culture-based method is time-consuming and requires specific culture conditions for *P. larvae* and *M. plutonius* strains. To avoid these limitations associated with culture methods, molecular diagnostic techniques such as PCR and qPCR are being used, which can be easily replicated and can detect more than one target in one reaction using specific primer sets. However, these methods detect both viable and dead bacteria and are indistinguishable. Therefore, in this study, we used propidium monoazide (PMA), a DNA-binding dye that inhibits DNA amplification from dead cells, allowing detection and quantification of only viable cells. The study aimed to evaluate PMA-qPCR as an alternative to culture for enumerating viable cells of AFB and EFB pathogens. First, we tested the effectiveness of PMA-qPCR to detect and quantify viable *P. larvae* and *M. plutonius* in cell cultures. Then, we used the same method to detect and quantify viable *P. larvae* on AFB-infected honeycomb samples before and after ozone treatment. Overall, we demonstrated that PMA-qPCR can be utilized for the detection of viable AFB and EFB pathogens in cultures and hive samples, which will aid in initiating appropriate treatment before the spread, as well as in evaluating the effectiveness of any treatment methods employed.

In our study, we used three different cell dilutions at concentrations of 10^8^, 10^6^, and 10^4^ CFU/mL for *P. larvae*, and 10^6^, 10^4^, and 10^2^ CFU/mL for *M. plutonius* to determine pathogen detection limits using PMA-qPCR. Cells were heat-treated to obtain dead cells for PMA treatments. Both viable and dead cells were treated with 25uM PMAxx, followed by qPCR. The qPCR signals were detected not only from PMA-treated viable cells but also from PMA-treated dead cells (Supplementary Figure 1); however, the Ct values of PMA-treated dead cells were considerably higher than those of PMA-treated viable cells, distinguishing live from the dead cells (Figure 3). There are several possible explanations for the amplification of dead cells treated with PMA. It could be that there were remaining dead, but intact cells with not enough cell membrane disturbance from the heat-treatment, which may have hindered the PMA from entering the cells and binding to their DNA. For our study, we prepared dead cells by heating 1 mL of each cell dilution at 95-120° C for 20 minutes for *P. larvae* strains and 30 mins for *M. plutonius* strains, followed by checking cell viability by plating 100 µL of each dilution on their respective media plates and incubating for 3 days for *P. larvae* and 6-7 days for *M. plutonius* strains. Cell death was confirmed by plating assay (Table 1). Similar methods were practiced in other studies as well (Lv et al., 2020) (Forsgren et al., 2008). The other possible reason could be that the amount of bacterial concentrations might be too high for the PMA to work, leaving dead cells without sufficient crosslinked PMA. There are some studies that reported complete inhibition of the qPCR signals from the dead *Campylobacter* cells (Josefsen et al., 2010), whereas others failed to completely inhibit the signal of dead *Campylobacter* cells when using a similar PMA qPCR approach (Pacholewicz et al., 2013). Some studies used two rounds of PMA treatments to improve the efficiency of the inhibition of dead cells (Okada et al., 2022).

For our study, we used SYBR-green master mix for the qPCR assay. Some studies reported qPCR assay for the detection and quantification of *P. larvae* spores in hive debris based on SYBR technology have limitations of being less specific and sensitive when compared with TaqMan-based probes, and cause difficulty in distinguishing highly similar gene amplicons from closely related species (Rossi et al., 2018; Tajadini et al., 2014). These specificity issues make it less appropriate for reliable quantification (Kušar et al., 2021; Schwendener et al., 2025). However, our melt curves showed only one amplicon type. There are no articles in the literature reporting that PMA-qPCR with SYBR-green was not reliable for detecting and quantifying *P. larvae* or *M. plutonius*. However, for other pathogens, a paper reported that the intercalating dye SYBR Green can detect viable *Campylobacter* cells among dead cells if present at a concentration of 6 log colony-forming units (CFUs/mL) (Lv et al., 2020), and TaqMan probe-based qPCR showed higher specificity than SYBR Green-based qPCR in the detection of *Campylobacter* spp. (Botteldoorn et al., 2008). While for *Brucella* sp., a study reported that SYBR-green PMA-qPCR effectively detected and quantified viable pathogen (Zhang et al., 2020).

Our results showed that there were lower *P. larvae* genome copies in the PMA-treated comb than in the PMA-untreated comb samples, before ozone treatment (Figure 4), suggesting that the viable *P. larvae* cells in the comb samples were significantly lower than what the qPCR results showed without PMA-treatment. This result suggested that PMA-qPCR can be used on hive samples to differentiate between viable and dead pathogens and to quantify viable *P. larvae* in the samples. To the best of our knowledge, there have been no reports using PMA-qPCR to quantify AFB and EFB pathogens from the hive samples. However, several studies have used this method to quantify other pathogens. For example, it has reported that the use of PMA increased the ability to resolve the impact of antibiotic therapy on *Pseudomonas aerugionasa* load in cystic fibrosis respiratory samples (Rogers et al., 2010), to determine viable cells of inter-species biofilm of *Candida albicans*-*Staphylococcus aureus* (Kendra et al., 2024), and to accurately detect live cells of *Vibrio vulnificus* in clinical samples to enhance public health safety and to improve the emergency response level for *V. vulnificus* infection (Hu et al., 2022). It has been used in the agri-food production system for the detection and quantification of viable but non-culturable *Campylobacter jejuni* in poultry products, a foodborne pathogen causing gastrointestinal infections in humans (Lv et al., 2020). It has also been used in environmental samples, wastewater treatment (Heise et al., 2016; Nocker et al., 2007), to detect viable *Escherichia coli* in agricultural soil contaminated by manure, an important source for the transmission of foodborne pathogens (Fu et al., 2020).

After an hour of ozone exposure, the genome copies of *P. larvae* in both the PMA-treated and the PMA-untreated comb samples decreased, and the difference between them was lower. This suggests that the decrease in genome copies after 1 hour of ozone exposure may be due to the degradation of extracellular DNA. Similar results have been reported, showing that ozone can be used as a fumigant and is detrimental or lethal to *P. larvae* (James, 2011), including other pests and pathogens (Rangel et al., 2021). A study by Torlak & Işik (2018) evaluated the efficacy of two ozone concentrations (9.8 and 17.1 mg/L) for up to 120 minutes at room temperature in inactivating the spore cocktail of three *P. larvae* strains on wooden (pinewood stick) and plastic hive materials (PVC sticks). After ozonation, spore reduction was significantly higher on PVC sticks than on pinewood sticks, suggesting that ozone treatment is more promising for decontaminating plastic hives. Wooden materials have porous surfaces that spores can penetrate and become embedded in the cavities, making decontamination difficult. One important point to note is that the ozone treatment at 17.1 mg/L for 120 minutes yielded over 4 log reduction in the counts of the spores on PVC sticks, but didn’t eliminate the spores completely. This is what we have observed in our studies (Figure 4). We observed lower genome copies; however, the Ct values of PMA-treated and the PMA-untreated samples, even after ozone exposure, were significantly different (Figure 4). This suggests that some viable cells may have survived the ozone exposure and appeared in the qPCR without PMA, as well as in the samples after PMA treatment. It could be the ozone concentration wasn’t high enough or the 1-hour exposure time was too short, which might hinder the elimination of the entire spores from the samples. Another study reported that *P. larva*e were effectively sterilized or killed with ozone at a higher concentration of 8,560 mg O_3_/m^3^ and longer exposure periods of 3 days, under conditions with high temperature at 50°C and relative humidity higher than 75% (James, 2011). Therefore, further experiments are warranted using similar conditions.

Here we used AFB-infected hive samples and did not have access to any naturally infected EFB samples. Therefore, it is worth testing the EFB-infected environmental samples using PMA-qPCR for detecting and quantifying the pathogen.

## CONCLUSION

In this study, we assessed the performance of PMA-qPCR for detecting and quantifying viable *Paenibacillus larvae* and *Melissococcus plutonius*, first in pure cultures and subsequently in environmentally relevant honeycomb samples before and after ozone treatment. PMA treatment substantially reduced the erroneous detection of dead cells of both pathogens in culture and hive samples by several orders of magnitude. To our knowledge, this is the first study to apply PMA to detect AFB and EFB pathogens in honeybees. This approach has potential utility for AFB and EFB risk assessment and for evaluating the efficacy of pathogen control strategies. However, PMA did not completely suppress PCR amplification from dead cells, indicating that further optimization of treatment conditions is needed for precise quantification of viable pathogens. In addition, further experiments addressing current limitations and optimizing PMA protocols across different AFB and EFB strains will enhance the reliability and applicability of PMA-based molecular methods as practical and quantitative tools for investigating honeybee pathogen dynamics in apiaries.

## ACKNOWLEDGEMENT

We thank Peter Daniel Fowler and Meghan Milbrath (Michigan State University) for providing the *Melissococcus plutonius* strain (19-18-9) used in this study. We thank Dr. Gary Chastagner (Washington State University) for providing us with the ozone chamber and Bri Price and Joey Rosario for collection and treatment of the AFB infected comb samples. We thank Lucelia De Moura Pereira and Yva Eline (University of Idaho) for their support in the lab.

