## supplementary_figures for "Propidium monoazide (PMA) with quantitative PCR to detect and quantify viable *Paenibacillus larvae* and *Melissococcus plutonius* in honeybee hive samples"

**Supplemental Data** – The supplied Excel file contains partially processed data. Each sheet contains data from a particular analysis. Sheet "cell_culture_qpcr" contains the Ct-values for the cell culture-based testing of PMA. The column, "rel_conc" indicates the amount of dilution, "obs_rel_conc" shows $2^{\Delta Ct}$ relative to undiluted replicates. "Perc_err" values were calculated according to (Expected Ct – Observed Ct / Expected Ct) * 100

### Supplementary Figures

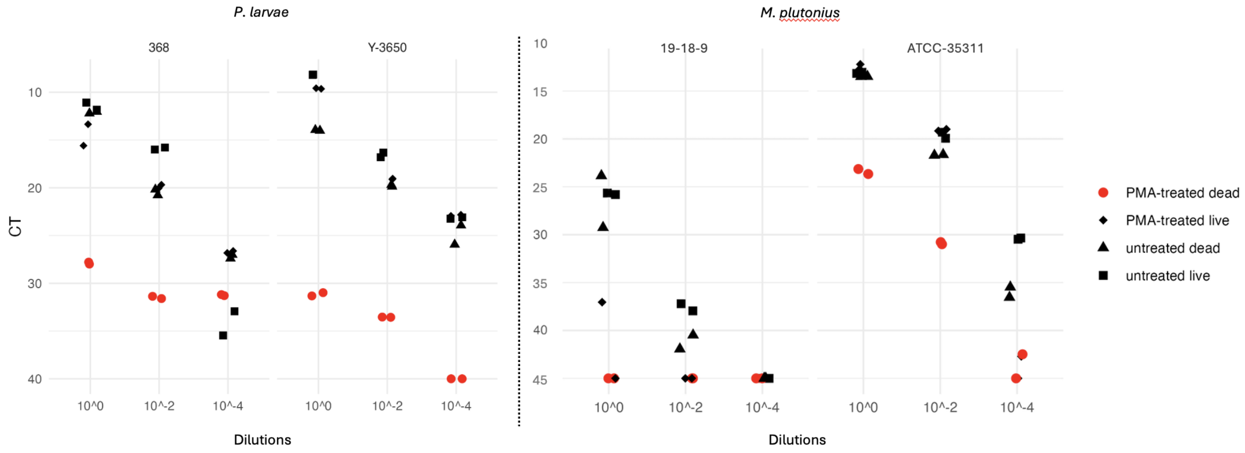


**Supplementary figure 1.** PMA-qPCR measurement of heat-killed cell cultures of bee pathogens. The difference between PMA-treated and untreated dead cells is shown in the manuscript (Fig 3). The dilutions $10^0$, $10^{-2}$, $10^{-4}$ correspond to cell densities of $10^8$, $10^6$, and $10^4$ for *P. larvae* and $10^6$, $10^4$, $10^2$ CFU/mL for *M. plutonius*.

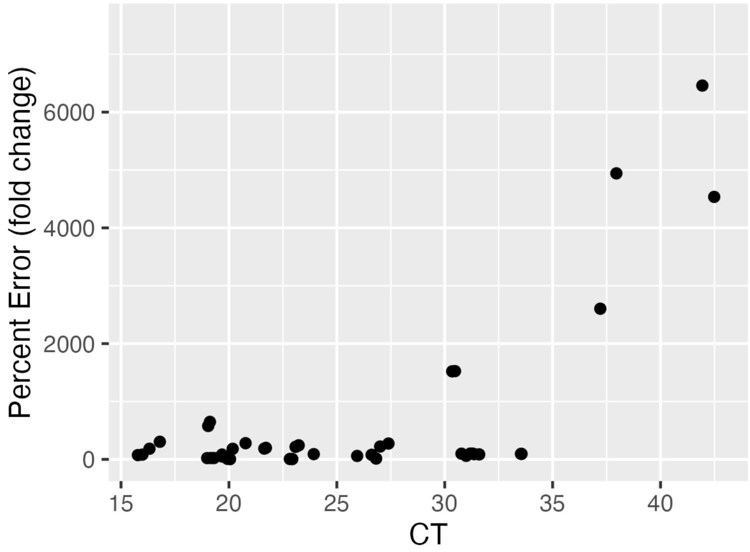


**Supplementary figure 2.** The calculation of error based on expected Ct values indicates that the qPCR assay becomes unreliable at higher Ct values. Percent error was calculated by calculating ΔCt from the $10^0$ dilution for each replicate. We then converted ΔCt to ΔGenomeCopy number using $2^{\Delta Ct}$. Percent error was calculated using the equation 100*((Expected copy number – Observed copy number) / Expected copy number), where expected is either a $10^{-2}$ or $10^{-4}$ dilution of the starting sample.

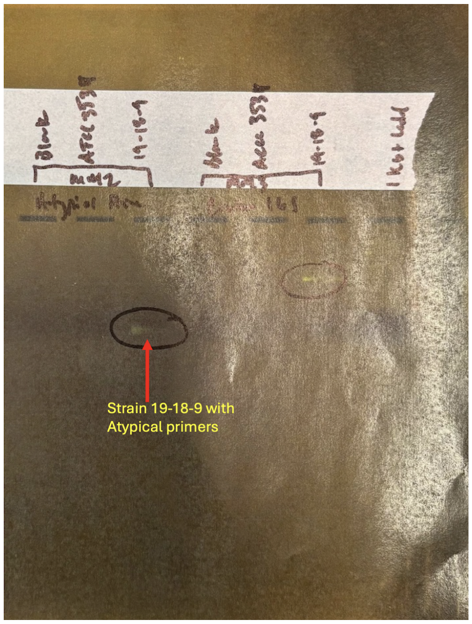


**Supplementary figure 3.** Strain 19-18-9 with atypical primers with dim bands when visualized on an agarose gel.
